# Uncovering the representational geometry of durations

**DOI:** 10.64898/2026.03.29.715088

**Authors:** Camille L. Grasso, Ladislas Nalborczyk, Virginie van Wassenhove

**Author notes:** Correspondence concerning this article should be addressed to Camille L. Grasso.

## Abstract

Is there a geometry of time in the human mind? A canonical measure of time in psychology is duration, a time interval quantifiable as a magnitude. Durations have been proposed to be arranged along a mental timeline: a unidimensional, linear, and spatialised representation of time. Here, we asked whether such a mental timeline is sufficient to account for the experience of duration. To address this, we tested the same participants in two experiments: a behavioural similarity judgment task, in which participants rated the similarity of duration pairs, and an electroencephalography (EEG) experiment in which they detected oddball durations. We used behavioural and EEG data to construct representational dissimilarity matrices, whose geometry was compared against theoretical models of duration organisation. Most variance in behavioural similarity judgements is explained by three latent dimensions, interpretable as: magnitude (monotonic ordering of durations), contextual encoding (distance to the geometric mean of the duration set), and a periodic component. These three dimensions are jointly consistent with a latent generalised helical model, which provided excellent fit to the behavioural data. Individual helical model parameters further correlated with endogenous neural oscillations measured during rest, suggesting that an individual’s duration space is partially shaped by intrinsic dynamics. The neural geometry was also found to be dynamic, unfolding in two successive stages: a strong logarithmic encoding of durations peaking around 150 ms after duration offset, followed by a spring-like geometry starting around 300 ms after offset. Together, these findings characterise the psychological and neural geometry of duration space.

## Introduction

All experiences unfold in time, making time a fundamental dimension of perception, cognition, and action. Characterising how the brain represents time is therefore a central challenge for cognitive neuroscience (e.g., 1, 2, 3, 4, 5, 6). However, the representational principles governing the coding of time intervals across scales, from subsecond to suprasecond durations, remain poorly understood. What, then, is the structure of the internal space in which durations are represented?

Several influential frameworks have proposed that time is spatialised along a mental timeline (e.g., 7, 8, 9). In many of these accounts, durations are conceived as magnitudes organised along a single internal dimension, with shorter and longer intervals ordered monotonically along a time axis (e.g., 10, 11, 12). This view captures important features of duration judgement but leaves unresolved the structure of duration representations themselves. More specifically, whether the psychological and neural representations of duration can be reduced to a single latent axis, or whether they are better characterised as a higher-dimensional geometry remains to be addressed.

A constructive way to address this question is to adopt a relational approach to the representation of durations: instead of asking how a given duration is represented in isolation, we can ask how durations are positioned relative to one another in terms of perceived similarity. This idea builds on the foundational insight that, even when direct correspondences between internal representations and external referents are difficult to establish, lawful structure may still be recovered from the relations among representations (*13*). Pairwise similarity judgements offer an empirical window onto the structure of mental representations, while multidimensional scaling (MDS) provides a formal tool for recovering the latent geometry that organises those judgements (e.g., 14, 15, 16, 17, 18, 19, 20).

This relational logic has been extended to neural data through representational similarity analysis (RSA) (*21*). In this framework, activity patterns elicited by each condition are treated as points in a high-dimensional space, and the pairwise dissimilarities between these patterns are summarised in a representational dissimilarity matrix (RDM). Crucially, because RDMs are defined over conditions rather than over specific sensors, neurons, or voxels, they provide a common representational format that can be compared across behavioural data, neural recordings, computational models, individuals, and species (*21*). RSA and related approaches have been applied across a broad range of cognitive domains, including perception (e.g., 22, 23), abstraction (e.g., 24, 25), decision-making (e.g., 26), and social cognition (e.g., 27, 28).

A key advantage of the relational framework is that the geometry of representational spaces can reveal the computational principles that structure a given cognitive domain. For instance, in colour perception, pairwise similarity judgements have long been known to recover an internal organisation closely related to the perceptual colour wheel (e.g., 29). Recent work has shown that the geometry derived from similarity judgements is sufficiently sensitive to capture subtle differences in colour experience across the visual field (e.g., 30), between groups of colour-neurotypical and colour-atypical observers, and between humans and large language models (e.g., 31, 32). At the neural level, multivariate analyses of magnetoencephalography (MEG) responses have revealed a dynamic colour geometry predictive of universal colour-naming patterns, illustrating how representational geometry can bridge neural activity and perceptual experience (*33*). Similar principles have also been identified in audition, where pitch perception is organised along a helical structure (e.g., 34, 35). In this helical structure, tones separated by an octave occupy analogous positions across successive turns. This geometry captures both monotonic variations in pitch height and periodic variations in chroma (e.g., 36, 37, 38, 34, 35). These examples show that similarity structure can reveal representational organisations that are not reducible to a single linear axis.

We therefore applied this approach to time perception and asked how many latent dimensions organise the human psychological space of durations. By "dimension", we refer to a latent organising axis within a psychological space, here termed the duration space. In this view, duration space is a representational geometry in which durations vary systematically, such that pairwise similarities between them can be approximated by distances in a low-dimensional space. These dimensions are not assumed to map one-to-one onto single neural variables (i.e., they do not imply isomorphism; 39, 40); rather, they provide a formal bridge between behavioural and neural representational structures.

We tackled this question by combining behavioural similarity judgements with electroencephalography (EEG) recordings within a common representational framework. In a first experimental session, participants rated the perceived similarity of all possible pairs of auditory durations. Behavioural data served the estimation of the geometry of duration representations in participants’ psychological space. In a second session, we recorded EEG while the same durations were presented to the same participants. We derived RDMs from EEG recordings and the multivariate similarity structure of evoked responses. Behavioural and neural RDMs were then compared with the predictions of theoretical models of duration organisation.

We hypothesised that the psychological space of durations would not be adequately captured by a single dimension, but would instead require at least two latent dimensions (*41*). Our findings went beyond this prediction. They suggest that at least three organising principles are needed to account for the relational structure of duration space. The first is a magnitude dimension, reflecting the monotonic ordering of durations, potentially in a compressive form consistent with scalar timing. The second is a contextual dimension, capturing the encoding of durations with respect to the centre of the experienced distribution. The third is a periodic dimension, which may reflect similarity relations induced by endogenous neural rhythms. At the neural level, geometry was found to be dynamic, unfolding in two successive stages: a strong logarithmic encoding of durations peaking around 150 ms after duration offset, followed by a spring-like geometry, converging with the behavioural structure, starting around 300 ms after duration offset.

## Results

Participants judged the perceived similarity of pairs of auditory durations and, in a separate EEG session, detected rare duration oddballs embedded in a continuous stream of the same durations (Fig. 1).

**Figure 1.**
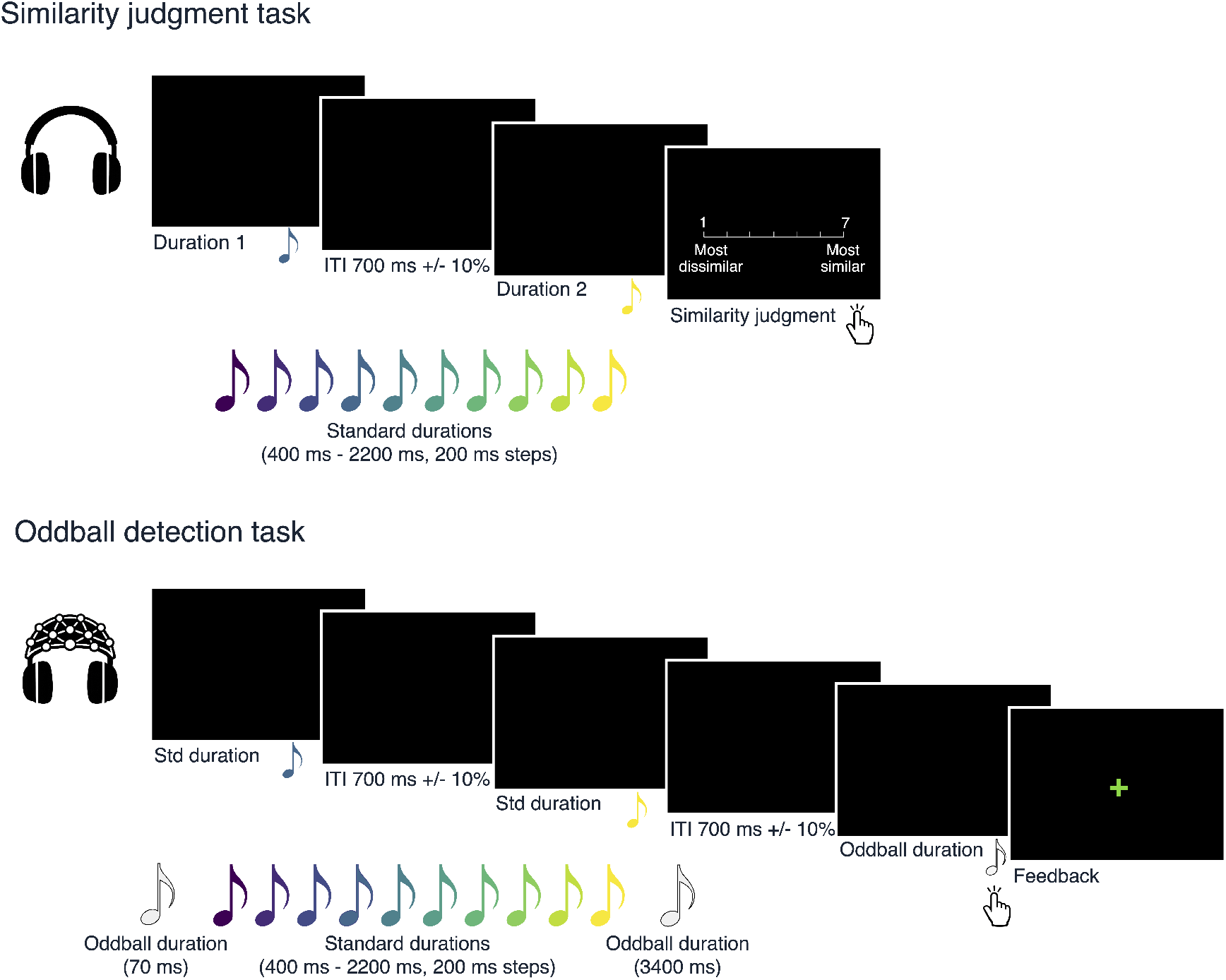
Overview of the experimental protocol. **(Top)** Similarity judgment task. On each trial, participants heard two tones in succession and judged how similar their duration felt on a 7-point Likert scale, from "most dissimilar" to "most similar". **(Bottom)** Oddball detection task. Participants listened to the same durations as in the behavioural experiment, but this time they were presented as a continuous stream with varying ITIs. Participants were tasked to detect rare duration oddballs, which were shorter or longer than all other standard durations. Responses were made by pressing the space bar, and visual feedback indicated correct detections and errors.

### Psychological duration space

We first asked whether participants’ similarity judgements revealed a coherent representational structure for duration, and if so, what its broad organisation looked like at the group level. Fig. 2A shows the average behavioural RDM at the group-level. As expected, the matrix displays a clear diagonal structure: pairs of similar objective durations were judged as more similar. The matrix also reveals an apparent decrease in discriminability with increasing duration: dissimilarities in the upper-right corner (longer durations) are generally lower than those in the lower-left corner (shorter durations). Interestingly, RDMs at the participant-level exhibited substantial inter-individual variability in both structure and noise (fig. S1), such that some fine-grained patterns were attenuated or lost when averaging across participants. This underscores the importance of analysing representational structure at the level of individual participants.

**Figure 2.**
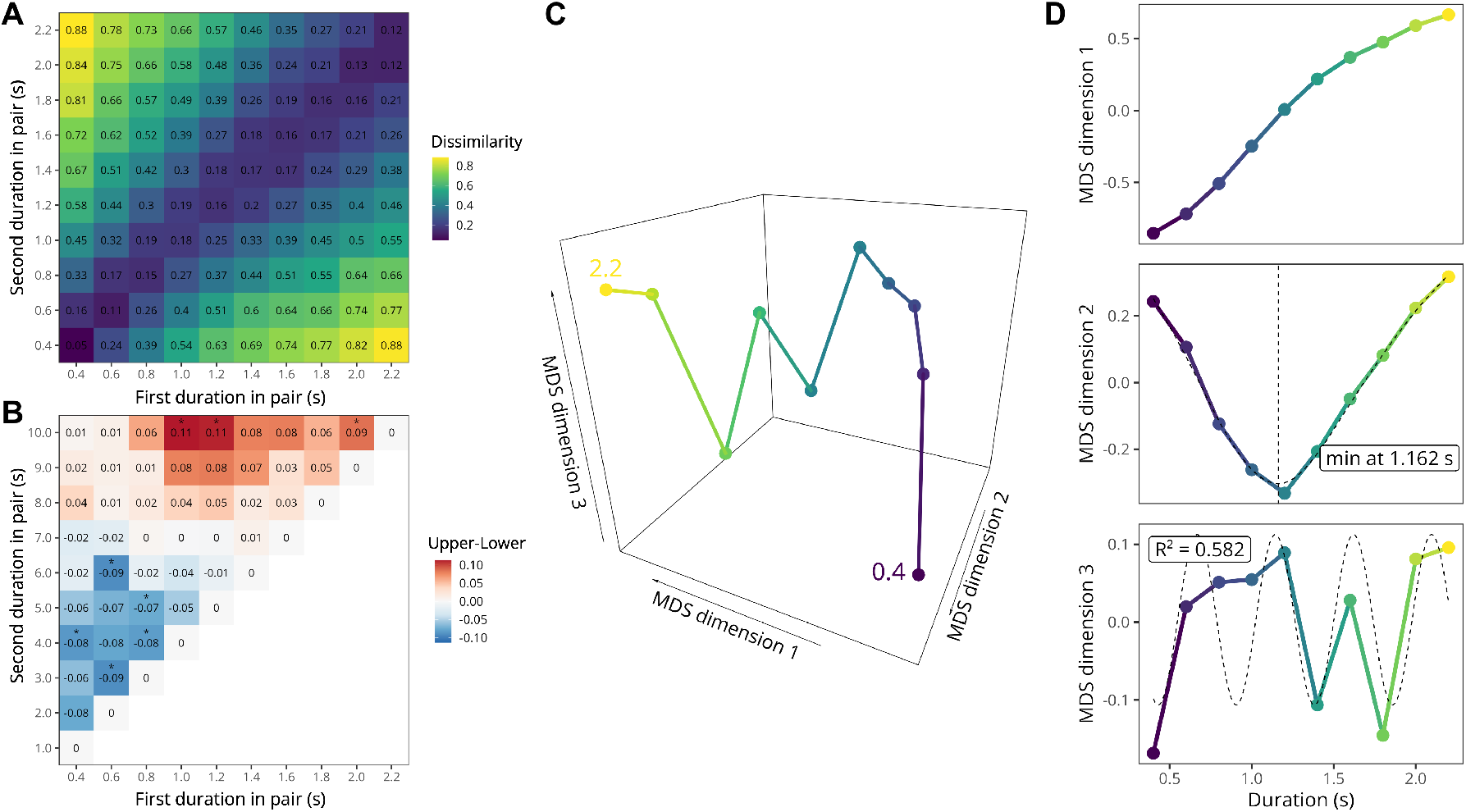
Geometry of the duration space (behaviour). **(A)** Empirical dissimilarity matrix derived from duration similarity judgements averaged at the group-level. Each cell shows the mean dissimilarity assigned to a pair of durations (s). Lower values (blue) indicate that two durations were perceived as more similar, whereas higher values (yellow) indicate that they were perceived as more dissimilar. The matrix shows a clear diagonal organisation, with dissimilarity increasing as the difference between durations grows. **(B)** Group-level asymmetry matrix, computed as A = D − D^T^. This panel highlights directional deviations from perfect symmetry in pairwise judgements. Red cells indicate pairs for which the dissimilarity was greater in one presentation order than in the reverse order (positive asymmetry), whereas blue cells indicate the opposite pattern (negative asymmetry). Values close to zero (white) indicate little or no order-dependent asymmetry. Stars indicate pairs of durations with statistically different order asymmetries (BF10s > 3). **(C)** Three-dimensional MDS solution fitted to the average empirical dissimilarity matrix. Each point corresponds to one duration. Neighbouring points represent durations judged as more similar. The embedding reveals a structured trajectory across durations rather than a simple linear arrangement. **(D)** Coordinates of the three MDS dimensions plotted as a function of objective duration. The first dimension varies monotonically with duration, suggesting that it captures the main ordinal progression from short to long intervals. The second dimension shows a U-shaped profile with a minimum around 1.16 s, indicating sensitivity to distance from a central tendency. The third dimension shows a periodic-like modulation across durations (which can be described by a sinusoidal function, represented by the dashed curve), consistent with an additional oscillatory or cyclic component in the representational geometry.

The asymmetry matrix, computed as *A* = *D* − *D^T^*, reveals a striking and systematic order effect (Fig. 2B). For duration pairs shorter than the average duration (1300 ms), the first-presented duration tended to be judged as longer than when the same duration appeared second. For pairs longer than the average of the tested distribution, the pattern reversed. Because these asymmetries reflect task-induced, order-dependent biases, rather than intrinsic properties of duration representation, all subsequent analyses focus on order-invariant structure. To this end, we symmetrised each participant’s RDM prior to MDS and model comparison.

To uncover the geometry of the duration space at the group-level, we first aligned individual MDS solutions to a common coordinate frame using generalised Procrustes analysis (*42*). Then, we averaged the resulting 3D coordinates across individuals (Fig. 2C). A comparison with INDSCAL; a hierarchical MDS procedure; yielded qualitatively similar embeddings. A scree plot analysis of stress values revealed an elbow at three dimensions, beyond which higher-dimensional solutions provided only marginal improvements (fig. S2). We therefore retained the 3D MDS solution for all subsequent analyses.

The resulting 3D embeddings reveal a clear non-linear organisation of subjective durations (Fig. 2C), resembling a corkscrew or spring-like trajectory predominantly oriented along the objective-duration axis. This first dimension captures a monotonic ordering of durations by magnitude, scaling subjective duration with physical duration. The second dimension appears to encode the eccentricity of a duration relative to the geometric mean of the set of durations. The third dimension captures a periodic component, with clear peaks and valleys.

Fig. 2D also shows the 1D projections of the 3D group-level MDS embeddings in relation to objective duration. This figure confirms that the first MDS dimension arranges durations in a monotonic fashion. The second MDS dimension arranges durations according to their "distance" to a value (≈ 1.162 s) very close to the geometric mean of the durations set (≈ 1.151 s). The third MDS dimension arranges durations in a periodic pattern that was adequately described by a sinusoidal function (optimal period of ≈ 476 ms, *R*^2^ ≈ 0.582).

We next tested whether the higher MDS dimensions could be explained by a classical horse-shoe artifact (*43, 44*). Specifically, we asked whether any higher-dimensional structure remained reliable across participants after removing polynomial functions of Dimension 1. To this end, we residualised MDS Dimensions 2 and 3 with respect to a third-degree polynomial of Dimension 1 and computed pairwise distances in the resulting residual two-dimensional space. Across random participant split-halves, the residual distance structure showed positive reliability, median (r = .33), 95% interval ([-.04, .70]). This reliability was substantially greater than a permutation baseline in which duration-pair labels were shuffled, median permuted (r = −.01). Observed reliability exceeded the permuted value in 88.7% of splits, Wilcoxon signed-rank test, (*p* < .001). Thus, although Dimension 2 contained a clear horseshoe-like component, the higher-dimensional configuration was not fully reducible to polynomial transformations of Dimension 1.

### Model comparison

Given the large degree of inter-individual variability in behavioural similarity judgements (cf. fig. S1), we performed the subsequent analyses at the participant level. Fig. 3A shows the results of the model comparison procedure for one exemplary participant. This figure shows that, overall, all theoretical RDMs provided a good fit to the empirical RDM (all Spearman’s *ρ >* 0.7) with similar looking patterns.

**Figure 3.**
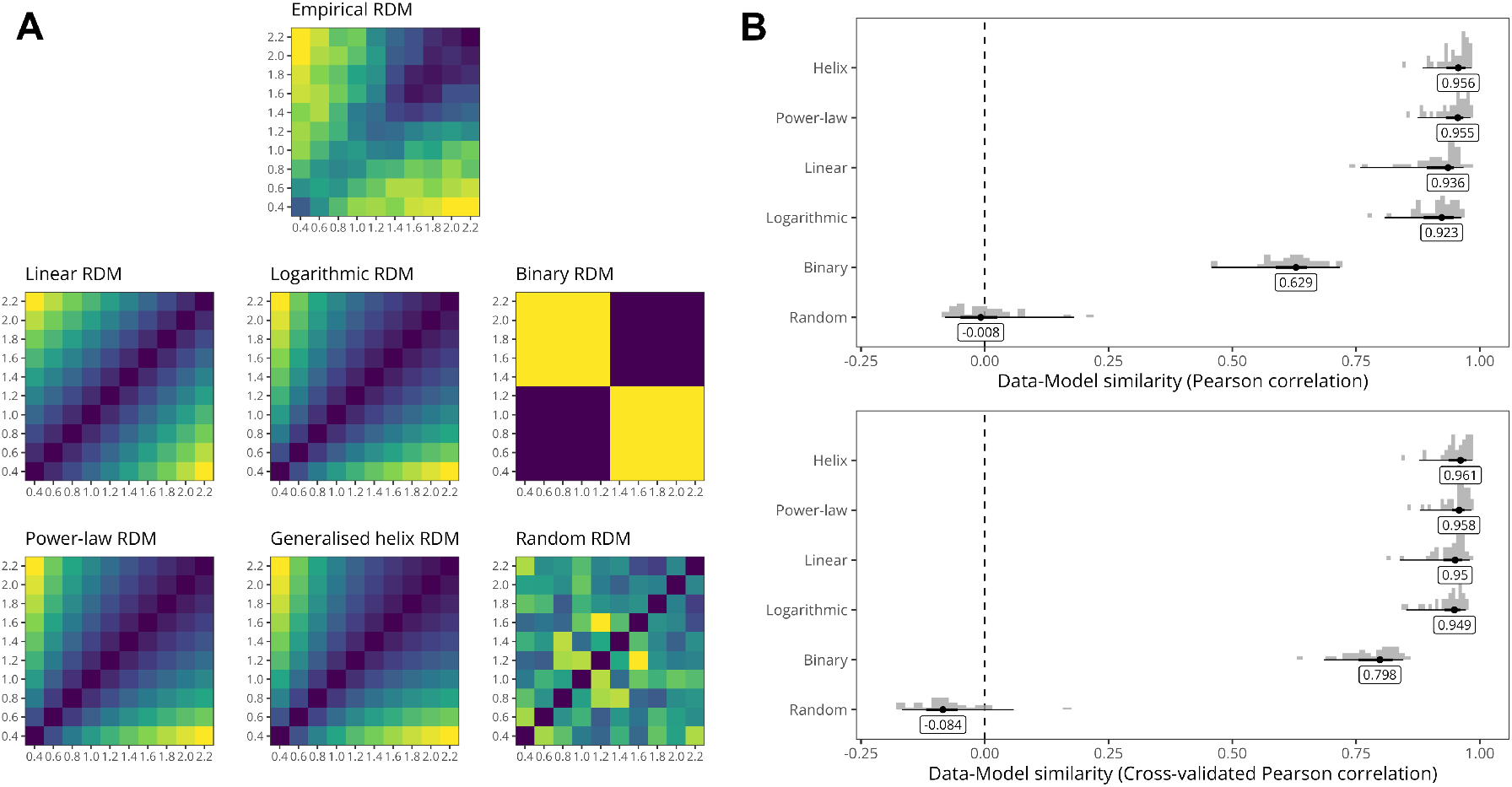
Behavioural RSA results. **(A)** RSA results for one exemplary participant. The top row shows the average symmetrised empirical RDM (i.e., subjective similarity ratings) whereas the second and third rows show the model RDMs. **(B)** Group-level results of the model comparison. Top panel shows the raw Spearman correlation for each model, whereas the bottom panel shows the cross-validated score. In both panels, models are vertically arranged by their median similarity to the empirical RDM. Densities show the distribution of similarities across participants. The dots and error bars show the median along with the 50 and 95 % central quantiles of this distribution.

Fig. 3B presents the group-level RSA results, showing the distribution of raw optimised correlation values (top row) and cross-validated correlation values (bottom row) for each candidate model. Models are ordered vertically according to their performance, indexed by the median cross-validated correlation. Overall, two models consistently provided the best fit to the empirical RDMs: the Helix and Power models. Importantly, the bottom panel indicates that cross-validation preserved the relative ordering of models, suggesting that model ranking was not driven by overfitting. However, cross-validation attenuated the differences between models, notably reducing the performance gap between the optimised models (i.e., Helix and Power) and the non-optimised models (Linear and Log models).

This pattern was confirmed by the Bayesian multilevel analysis of the fold-level cross-validated scores. Posterior pairwise comparisons between models were computed from the posterior distribution of expected CV correlations, after back-transforming estimates from the Fisher *z* scale to the correlation scale. The Helix and Power models showed virtually indistinguishable performance: the posterior distribution of the Helix–Power difference was centred close to zero, with a median difference of 0.002, and almost all posterior mass falling inside the ROPE (99.97%; fig. S3). In contrast, both optimised models showed higher expected cross-validated correlations than the Linear and Log models. The Helix model outperformed the Linear model (median difference = 0.014; 100% posterior mass above zero; 13.27% inside the ROPE) and the Log model (median difference = 0.018; 100% posterior mass above zero; 0.82% inside the ROPE). Similarly, the Power model outperformed the Linear model (median difference = 0.012; 100% posterior mass above zero; 25.54% inside the ROPE) and the Log model (median difference = 0.016; 100% posterior mass above zero; 2.37% inside the ROPE). By contrast, the Linear and Log models were not meaningfully distinguishable, with a small median difference (0.004) and most posterior mass inside the ROPE (91.14%). Together, these results indicate that the Helix and Power models provided the best account of behavioural representational geometry among the tested models, while the available evidence did not support a practically meaningful difference between these two models.

### Event-related potentials

Having characterised the geometry of duration space at the psychological level, we next examined its neural geometry. We focused on activity time-locked to duration offset, as this marks the point at which the full temporal interval has become available to the system. This alignment is therefore the most direct for testing neural signatures of represented duration. All analyses reported below were performed on offset-locked data. Fig. 4A shows the group-level scalp distribution of the *t*-values associated with the slope estimates from the rank linear regression, sampled over the 600 ms following stimulus offset. The effect was robust and temporally sustained across this post-offset window. Its topography showed negative *t*-values over fronto-central sensors (blue) and positive *t*-values over parieto-occipital sensors (red), revealing a clear anterior–posterior polarity.

**Figure 4.**
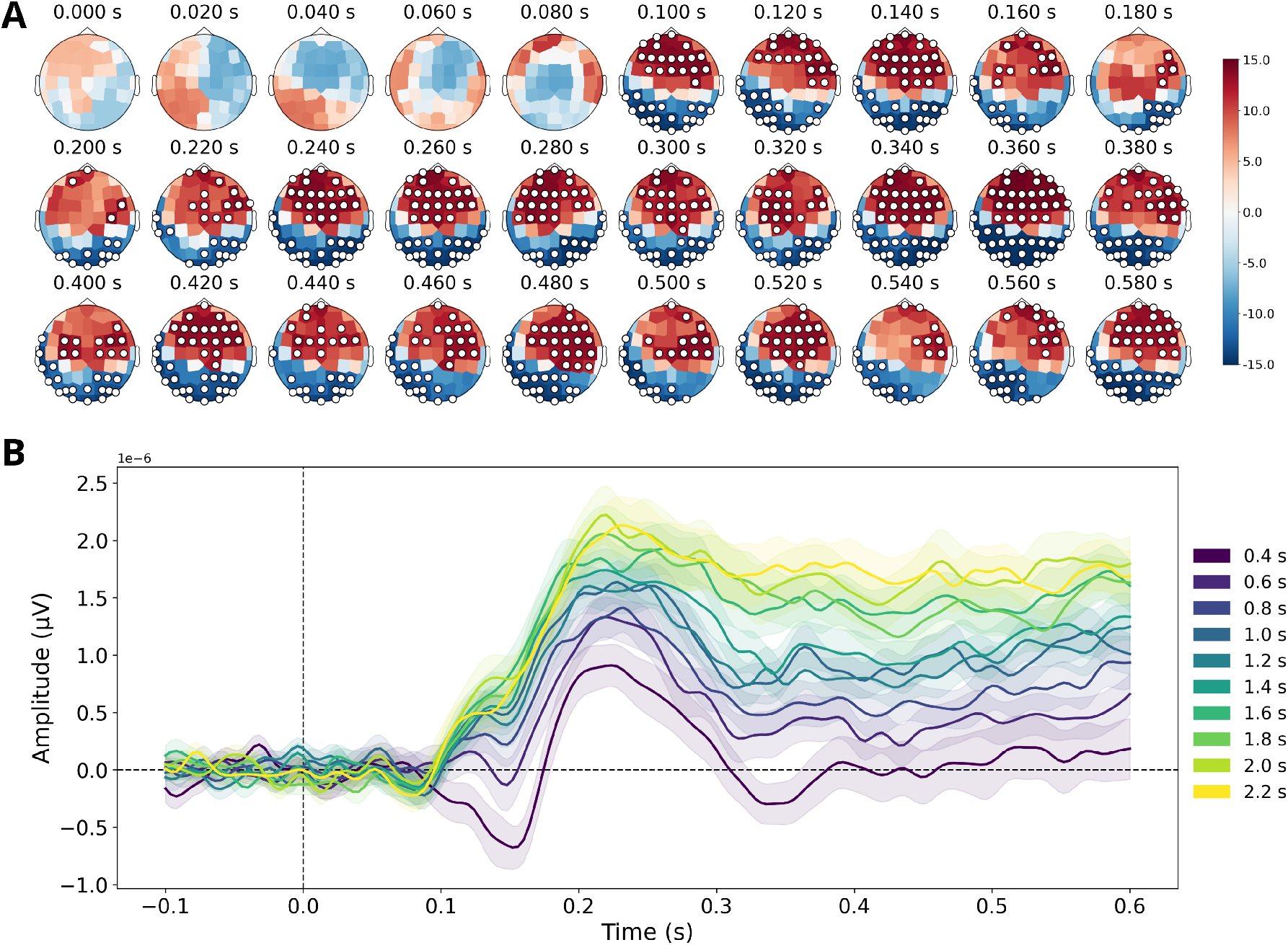
Event-related potentials at duration offset. **(A)** Group-level t-values of the slopes as estimated by the rank linear regression at duration offset. Spatio-temporal clusters (indicated by labelled sensors) were determined via spatio-temporal cluster-based permutation tests with TFCE and a significance threshold α = .05. **(B)** Group-level average ERP computed within the fronto-central cluster. Each coloured line represents the average EEG activity within this cluster for a given duration. The ribbon spans an interval of +/- 1 standard error of the mean.

Fig. 4B shows the group-level average ERP time-locked to stimulus offset for each duration, averaged over the fronto-central cluster of electrodes identified by the cluster-based permutation test (shaded areas indicate variability across participants). Shortly after off-set, the evoked response diverges as a function of duration. A pronounced positive-going deflection emerges around ≈ 120 − 150 ms, peaks around ≈ 180 − 220 ms, and is followed by a sustained positivity extending through ≈ 600 ms. Crucially, both the peak amplitude and the later sustained activity scale monotonically with duration: longer durations (green/yellow traces) show the largest positive amplitudes, whereas shorter durations (purple/blue traces) remain smaller and closer to baseline.

### EEG representational structure through time

Having established the behavioural geometry of duration judgments, we next asked whether a comparable structure could be tracked in neural activity, and if so, when it emerges over time. To address this question, we examined the group-level average EEG RDMs across successive post-offset time windows, together with their associated 2D MDS embeddings (Fig. 5). This analysis revealed a clear temporal transition from an initially weak and poorly differentiated geometry to a progressively more organised representational structure. In the earliest windows (0-120 ms), the embeddings remained highly compact, with durations tightly clustered and little systematic separation between neighbouring values, consistent with the relatively undifferentiated RDMs observed at the same timepoints. From around 160-200 ms onward, however, the representational space expanded markedly and began to show a clear ordering with duration: shorter intervals (e.g., 400-800 ms) tended to occupy one side of the space, whereas longer intervals (e.g., 1800-2200 ms) progressively separated toward the opposite side. This change was mirrored in the RDMs, which became increasingly banded, with lower dissimilarities concentrated near the diagonal and larger dissimilarities emerging for pairs farther apart in duration. Thus, by about 200 ms after offset, EEG activity no longer appeared to reflect only a weak or generic post-stimulus response, but rather a structured representation preserving relational information between durations. Importantly, however, this emerging organisation was not exhausted by a simple magnitude-like ordering, and the subsequent analyses suggest that additional structuring principles also contributed to the geometry observed in this time range.

**Figure 5.**
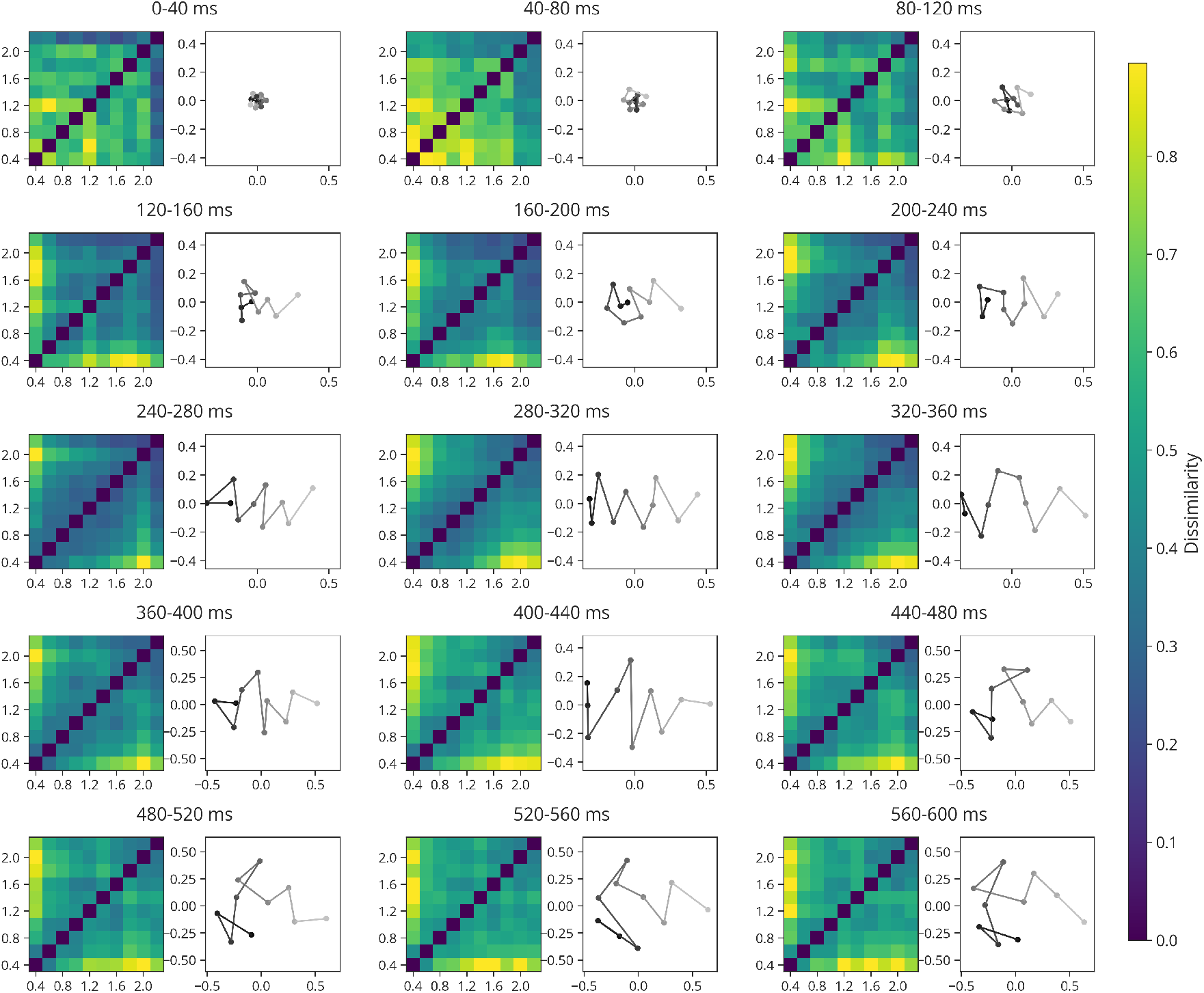
Dynamics of the neural geometry of duration space at stimulus offset. For each 40-ms time window from stimulus offset to 600 ms after offset, the left panel displays the group-average EEG representational dissimilarity matrix (RDM), in which each cell indicates the dissimilarity between the neural patterns evoked by a pair of stimulus durations. Lower values (blue) indicate more similar neural representations, whereas higher values (yellow) indicate more distinct representations. The right panel shows the corresponding 2D MDS solution for the same RDM, providing a geometric visualisation of the representational structure: durations plotted closer together were represented more similarly in the EEG signal, whereas durations plotted farther apart were represented more distinctly. Together, these panels illustrate how the geometry of duration space evolves over time following stimulus offset.

Crucially, in the 160-240 ms window, the RDMs exhibited an additional regularity that was not captured by a simple distance-from-diagonal gradient: dissimilarities appeared to depend not only on the absolute difference between two durations, but also on whether those durations occupied a similar position relative to the nearest oddball anchor (i.e., 70 ms or 3400 ms). In other words, durations that were similarly "regular" with respect to these anchors - that is, that had a comparable temporal distance to the closest odd duration - tended to evoke similar multivariate EEG patterns. This observation is important because it suggests that the neural geometry was already reflecting more than a straightforward short-to-long ordering. Instead, the representation appeared sensitive to a second-order property of the duration set: the relational status of each duration within the broader temporal context defined by the oddball anchors.

From approximately 200 ms onward, the embeddings showed a visually periodic component superimposed on the global short-to-long ordering, suggesting that the representational geometry was not strictly one-dimensional. Although the overall ordering of durations remained broadly preserved, the trajectory linking successive durations repeatedly bent back and forth through the space, revealing recurrent structure beyond monotonic scaling. This pattern is compatible with a second, cyclic organising principle layered on top of the magnitude-like axis, such as an oscillatory or rhythmic component in similarity space.

Taken together, these observations suggest that the post-offset neural representation of duration is not adequately described as a simple linear magnitude code. Rather than collapsing onto a single magnitude axis, the neural representation of duration appeared to combine magnitude, anchor-relative, and periodic structure.

### Representational similarity analysis

The MDS and RDM visualisations suggested that post-offset EEG activity progressively acquires a structured geometry, combining an overall magnitude-like ordering with additional relational and possibly periodic structure (Fig. 5). We next asked whether these qualitative impressions could be formalised using explicit theoretical models of duration representation. To do so, we used representational similarity analysis (RSA) and commonality analysis to quantify, at each timepoint, how much variance in the EEG RDMs could be explained by a set of candidate theoretical RDMs, both jointly and uniquely.

Fig. 6 shows the results of these analyses. We first asked whether the emerging post-offset neural geometry was sufficiently structured to be captured by formal representational models at all. Fig. 6A shows that the variance explained by the theoretical models increased sharply from approximately 100 ms after duration offset. The full model (dotted black line), containing all predictors, provided a ceiling estimate of performance, and accounted for approximately 40-50% of the total variance in the EEG RDMs, with peaks around 150 and 300 ms after offset. This indicates that the post-offset neural response was organised enough to be captured to a great extent by a principled model space.

**Figure 6.**
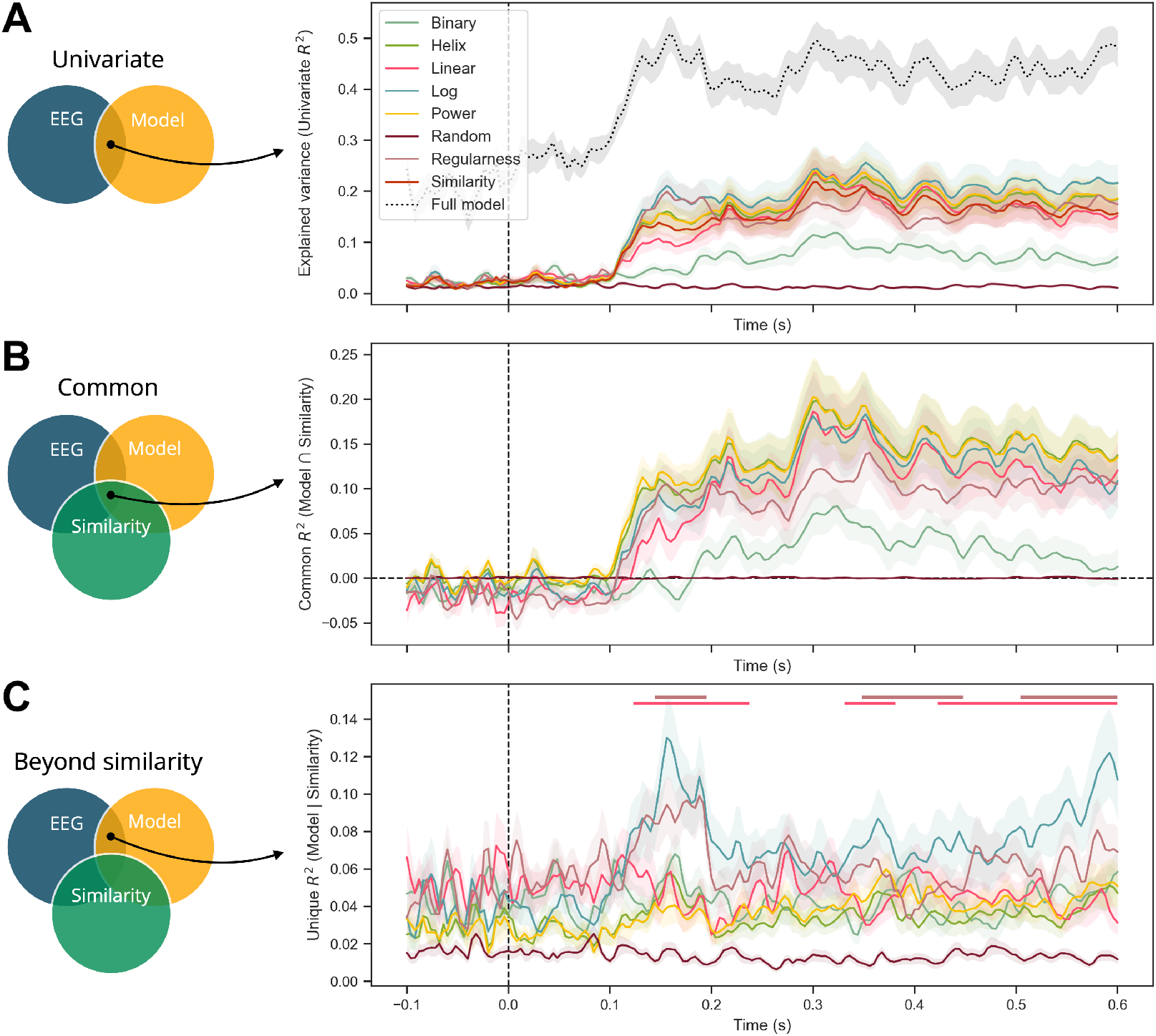
Group-level average EEG RSA and commonality results (+/- 1 SEM). **(A)** Explained variance for each predictor (as well as the full model). **(B)** Explained common variance between each model, the EEG RDM, and the behavioural similarity RDM. **(C)** Unique variance explained by each model beyond the behavioural similarity RDM. The brown horizontal bar indicates the timesteps where the Log model explained more variance than the Regularness model, whereas the pink horizontal bar indicates the timesteps where the Log model explained more variance than the Linear model (model-based posterior odds above 20).

Having established that several models explained the neural data, we next asked whether they did so because they tracked the same structure as the behavioural similarity, or because they captured additional properties of the neural representation. The upcoming commonality analyses clarify this point. The variance shared with the Similarity model (Fig. 6B) increased markedly after 100 ms and peaked around 300 ms. This was especially true for the best-performing structured models, for which a substantial part of their performance reflected representational structures also found in behavioural similarity judgements. This convergence is important because it links the post-offset neural geometry to the psychological geometry we characterised earlier. The two distinct levels of analysis (behaviour and EEG) recover a common representational organisation of duration.

Finally, we asked whether any model retained explanatory power beyond what could already be accounted for by behavioural similarity alone. Fig. 6C shows that two models retained substantial *unique* explanatory power beyond Similarity: the Log model showed the most pronounced transient boost around 150–200 ms, together with a later rise toward the end of the analysis window. Additionally, the Regularness model explained unique variance during the early post-offset period. These results suggest that the neural representation is not exhausted by the behavioural similarity structure alone. Rather, shortly after duration offset, the EEG signal appears to express additional representational principles; most notably logarithmic and anchor-relative regularity structure; that are only partially reflected in explicit similarity judgements.

Together, these results suggest a constructive sequence in the emergence of post-offset duration representations: first, time-resolved neural responses become sufficiently structured to be captured by formal models shortly after duration offset; second, a substantial part of this structure converges with the behavioural similarity geometry, especially around 300 ms; and third, some models; particularly Log and Regularness; capture additional neural variance around 150–200 ms that is not reducible to similarity judgements alone. Taken together, these findings indicate that the neural geometry of duration is both behaviourally grounded and richer than behavioural reports alone reveal.

### Task-related EEG correlates of duration representations

Having shown that duration is represented in the geometry of multivariate EEG patterns, we next asked whether this structure could also be detected in a simpler and classical electrophysiological signature of timing. We asked whether the post-offset P2 component (*45, 46, 47, 48, 49*) varied with duration in a way that recapitulated the magnitude-like and potentially non-monotonic structure revealed by the representational analyses. To address this, we quantified the relation between objective duration and the P2 amplitude and latency.

fig. S4 shows the group-level average P2 amplitude and P2 latency. The amplitude of P2 was found to grow monotonically with objective duration. The amplitude-duration relationship was reliably monotonic (*ρ* = 0.239, *t*(32) = 3.160, *p* = 0.0017), consistent with a progressive increase of P2 amplitude with longer intervals. In contrast, the latency of P2 exhibited a weaker and more heterogeneous association with duration. Although latencies tended to increase with duration in the group average, the Spearman’s correlation coefficient was smaller and did not reach significance (*ρ* = 0.102, *t*(32) = 1.470, *p* = 0.0757), suggesting that latency is not well captured by a simple monotonic trend and may instead reflect non-monotonic structure. Visual inspection of fig. S4 reveals local deviations from monotonicity around intermediate durations (1300 ms) and near the extreme durations. This pattern is reminiscent of the MDS pattern observed in the EEG RDMs (Fig. 5) as well as in the behavioural data (Fig. 2).

We found no significant correlations between participant-level mean P2 amplitude or latency, or oscillatory power at duration offset, and the parameter values of either the Helix or Power models.

### Endogenous oscillatory correlates of duration representations

We next asked whether the geometry of duration space was related to more stable, endogenous properties of the individual’s brain. To address this question, we tested whether inter-individual variation in the fitted parameters of the representational models was associated with resting-state oscillatory power, reasoning that intrinsic alpha and beta activity, both known for their implications in timing (*50,51,52,53,54*), might constrain how duration space is organised across participants.

Fig. 7 shows a positive correlation between the *β* helix parameter (governing exponential growth of the radius,see the Generalised helix model section in the Supplementary Materials) and both the alpha (Spearman’s *ρ* = 0.45, *p* = 0.019) and beta (Spearman’s *ρ* = 0.44, *p* = 0.012) power at rest. This suggests that participants with stronger alpha and beta power also had smaller (closer to 0) helix *β* parameter, indicating a slower exponential growth of the helix’s radius according to objective duration. In other words, participants with stronger alpha/beta power at rest had more spring-like helical structures, whereas participants with weaker alpha/beta power at rest had more conical helical structures, suggesting a stronger compression of larger durations. We found no other significant correlation between the Helix or Power model parameters and the endogenous oscillatory components we investigated.

**Figure 7.**
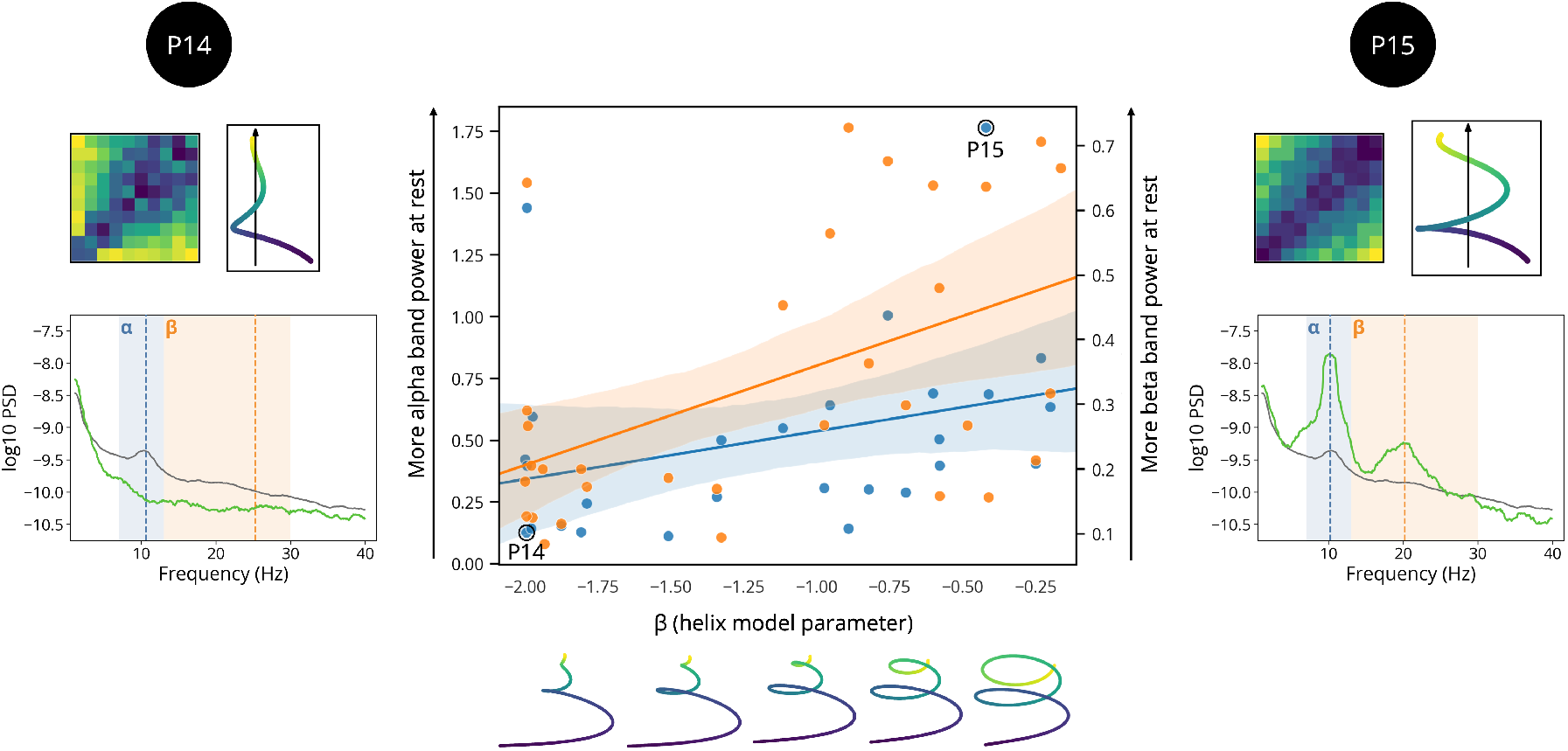
Relation between the helix β parameter and resting-state oscillatory power. Relation between participant-level estimates of the helix β parameter (x-axis) and resting-state power in the alpha band (blue, left y-axis) and beta band (orange, right y-axis). Higher resting-state alpha/beta power was associated with closer to zero helix β values, corresponding to more spring-like rather than cone-like helical structures, as illustrated by the 2D projections at the bottom. The lateral insets show two representative participants (P14 and P15): for each participant, the RDM represents the individual duration-similarity structure, the adjacent helix shows the fitted helical geometry, and the PSD panel shows the participant’s resting-state spectrum in green relative to the group-average spectrum in grey. In the PSD panels, alpha is highlighted in blue and beta in orange, with dashed vertical lines indicating the participant-specific spectral peaks used for power estimation.

The Bayesian regularised analysis provided a conservative assessment of the resting-state associations. The largest and most consistently positive effect was the association between Helix *β* and beta power (median standardised slope = 0.44, 95% CrI [−0.005, 0.89], *P* (*b >* 0) = .974), with little posterior mass inside the ROPE (2.3%). By contrast, the association between Helix *β* and alpha power, which was positive in the marginal robust correlation analysis, was strongly attenuated in the joint Student-*t* model (median standardised slope = 0.05, 95% CrI [−0.43, 0.54], *P* (*b >* 0) = .587). This indicates that the alpha-power association was not robust after accounting for shared variance with other EEG features. No regularised slope had a 95% credible interval entirely excluding zero. We therefore interpret these resting-state associations as exploratory rather than confirmatory, with the strongest remaining candidate association linking Helix *β* to resting-state beta power.

## Discussion

Duration is often treated as if it were represented along a single mental line running from short to long. The present results suggest a richer picture. Behavioural similarity judgements revealed a duration space structured not only by monotonic magnitude, but also by contextual position within the stimulus set and by a recurrent component not captured by standard scaling models. Time-resolved EEG analyses further showed that this structure emerges progressively after duration offset, with an early stage dominated by compressed and task-relevant coding and a later stage that more closely resembles behavioural similarity. These findings argue for a multidimensional and dynamic account of duration representation.

### Evidence for multidimensional structure in psychological duration space

The behavioural results provide the clearest evidence against a strictly one-dimensional account. First, as expected, durations were organised monotonically, with nearby durations judged as more similar than distant durations. This first dimension is consistent with a substantial literature suggesting that durations may be encoded monotonically, in a manner broadly analogous to other quantities. In the behavioural literature, this idea is often expressed through the notion of a mental timeline, according to which durations, like numbers, may be arranged along an ordered spatial axis, often from left to right (e.g., 12, 7, 9, 8). This proposal is supported by spatio-temporal congruency effects, whereby responses tend to be facilitated when short intervals or past-related concepts are mapped to the left side of space and long intervals or future-related concepts to the right (e.g., 55, 56, 57, 8, 58). More generally, interactions between spatial and temporal dimensions are consistent with magnitude-based accounts such as A Theory of Magnitude (ATOM), which posits partially overlapping neural resources for processing time, space, and number (e.g., 10, 11). Converging neuroimaging evidence has identified neuronal populations tuned to duration in cortical areas implicated timing and magnitude processing, including orderly topographic organisation of preferred durations in the supplementary motor area, described as chronomaps (*59*). Recent high-field fMRI work further suggests that duration coding follows a gradient from spatially dependent, monotonic encoding in early visual cortex to more abstract and spatially invariant coding in higher-level areas, culminating in the intraparietal cortex where spatial and temporal representations are closely intertwined (*60*). Together, these findings support the idea that one major component of duration representation is a monotonic magnitude dimension.

However, the behavioural space also contained a second dimension reflecting eccentricity relative to the centre of the stimulus range, with a minimum close to the geometric mean of the duration set. This pattern is consistent with the long-standing observation that duration judgements are context sensitive and gravitate toward the central tendency of the experienced distribution (e.g., 61, 62, 63). This pattern indicates that durations are not represented solely as absolute magnitudes, but relationally, with respect to the distribution in which they are embedded. Bayesian accounts formalise this by treating duration judgements as the combination of noisy sensory evidence with a learned prior over the stimulus distribution, naturally producing attraction toward the mean (*64, 65, 66*). Related modelling work further suggests that these effects are amplified when stimuli are randomised, or thought to be randomised (*67, 68, 69*). Recent latent-variable evidence converges with this interpretation, showing that temporal reproduction across sub- and suprasecond intervals is better captured by a general timing factor together with a second factor indexing regression toward the mean, rather than by a sharp discontinuity between interval-specific timing systems (*70*). In geometric terms, this implies that the structure of duration space reflects the position of each duration relative to the central tendency of the set, rather than the magnitude ordering only.

A third feature of the behavioural duration space was a periodic-like component super-imposed on the global duration ordering. This component is a novel and also a tentative aspect of the present findings. Importantly, we do not interpret it as evidence that duration is intrinsically "cyclic" in the same strong sense as pitch chroma. Unlike tones separated by an octave, which are perceived as highly similar despite differing in pitch height, durations that are far apart are clearly not experienced as equivalent or near-equivalent simply because they align at a similar "phase" of the fitted helix. Rather, the periodic component we observe should be understood as a weaker, secondary modulation of similarity relations layered onto the dominant monotonic organisation of duration. In other words, it indicates that pairwise duration similarity contains recurrent structure not captured by magnitude scaling or contextual compression alone. In that sense, the generalised helix should be understood as a parsimonious geometric model of the data: it provides a compact way of combining monotonic progression with recurrent variation in a single latent trajectory. Its good behavioural fit, preserved under cross-validation, suggests that this geometry captures genuine structure in participants’ judgements rather than merely overfitting idiosyncratic patterns. At the same time, the competitive performance of the Power model indicates that some aspects of the data can also be captured by monotonic nonlinearity alone. The main contribution of the helix is therefore not to replace all alternative models, but to make explicit that duration similarity may contain a recurrent component in addition to compression and ordering.

### A dynamic neuronal geometry of duration space

The EEG results further suggest that this representational geometry is assembled over time following duration offset. Shortly after offset, neural dissimilarity structure was best captured by logarithmic and regularness-based models, peaking around 150 ms. This early stage is compatible with a compressed coding of elapsed duration and with the extraction of task-relevant information about how each standard duration relates to the oddball anchors. By contrast, around 300 ms after offset, the neural RDMs became more strongly aligned with behavioural similarity and with the broader family of structured models, including the Linear, Power, and Helix models. This temporal sequence suggests that post-offset activity does not simply reflect a static readout of elapsed time, but a transformation in representational format: from an early stage dominated by compressed and task-contingent coding to a later, more stable geometry resembling the structure expressed in explicit similarity judgements. On this view, behavioural similarity may reflect a relatively late representational format that integrates multiple constraints rather than a direct readout of the earliest duration code.

These findings have several implications for theories of timing. First, they are broadly compatible with magnitude-based accounts in showing that duration is organised monotonically, both behaviourally and neurally (e.g., 7, 8, 9). Second, they extend scalar and psychophysical accounts (*71, 72*) by suggesting that compression is not merely a property of response variability, but is visible directly in the representational geometry itself. Third, they indicate that contextual structure, here expressed as distance to the geometric mean of the stimulus set, is not a secondary bias layered onto an otherwise metric representation, but may be one of the dimensions along which durations are encoded. This view is compatible with previous intracranial work in non-human primates showing that temporal context and prior expectations do not merely bias responses downstream, but reshape the geometry of population-level activity in frontal cortex during interval timing (*73, 74*). Finally, the periodic component raises the possibility that endogenous neural dynamics contribute to the geometry of duration space (as previously suggested by 41). The positive association between resting alpha/beta power and the fitted helix *β* parameter provides convergent, though indirect, support for the idea that stable oscillatory properties of the brain may shape how durations are represented (e.g., timing precision 75). One possible interpretation is that the periodic component reflects a trace of recurrent sampling mechanisms, broadly consistent with proposals that oscillatory dynamics can provide temporal coordinates for interval timing (e.g., 76, 77, 78). A non-exclusive, non-oscillatory account is that part of this profile reflects a gradual shift in the relative contribution of sensory-specific and modality-independent timing mechanisms near the subsecond range. However, such a transition would more naturally produce a local change or inflection, whereas the periodic-like modulation extended across the full 400–2200 ms duration set. This account may therefore contribute to the observed structure but is unlikely to explain the recurrent profile in full. However, the present data do not establish the mechanism underlying this component, and future work will be needed to determine whether it is genuinely oscillatory in origin, whether it depends on the specific range and spacing of the duration set, or whether it reflects a more general form of relational coding.

Comparable recurrent or rotational geometries have prominently been reported in motorcortical population activity during overt reaching movements (*79, 80*). Here, by contrast, the periodic-like structure emerged while participants monitored auditory durations, in analysed trials without overt responses and without rhythmic sensory or motor entrainment, suggesting that duration tracking can elicit recurrent representational structure without explicit movement production. More generally, the present findings fit with a broader shift from axis-based to manifold-based accounts of neural representation (e.g., 81, 82). In several domains, cognitive and perceptual variables are now understood as being encoded on low-dimensional manifolds embedded in higher-dimensional population activity, rather than along isolated scalar dimensions. In spatial cognition, for example, neural population activity has been shown to exhibit structured geometries including ring-like and toroidal manifolds, providing concrete examples of how continuous variables can be represented through recurrent low-dimensional organisation (e.g., 83, 84). Our results suggest that duration may likewise be represented in a geometry that combines monotonic progression with recurrent structure. At a more conceptual level, the cone-like expansion of the fitted generalised helix also resonates with broader attempts to think about temporality geometrically, including Bergson’s cone model of memory (*85*). These parallels remain heuristic rather than mechanistic, but they underscore a common point: once time is approached as a representational geometry, richer topologies than a simple line become both plausible and theoretically productive.

Several limitations should be acknowledged. First, the representational geometry was inferred from a specific set of auditory durations spanning 400 to 2200 ms in regular steps, so it remains unclear how far the recovered structure generalises across sensory modalities, temporal scales, and non-uniform sampling schemes. Although the similarity-judgement and oddball tasks differed in their broader temporal organisation (e.g., with different intertrial interval), both used a local inter-stimulus interval centred on 700 ms and jittered by ±10%. The cross-task convergence therefore argues against an explanation specific to the pairwise judgement structure, but does not exclude a contribution of the shared local inter-stimulus timing. Future studies should independently manipulate both duration sampling and ISI. Second, behavioural similarity judgements provide a powerful window onto representational structure, but they may also reflect decision strategies that are specific to the explicit comparison tasks. Third, the EEG analyses provide fine temporal resolution but limited spatial precision, preventing strong anatomical claims about where these geometries are implemented. Finally, the correlations with endogenous brain rhythms are correlational and therefore do not establish a causal contribution of endogenous rhythms to the geometry of duration space.

Despite these limitations, the present findings provide a step toward a geometric account of duration representation. Rather than supporting the idea of a single, one-dimensional, mental timeline, our results suggest that durations are organised in a richer representational space, structured behaviourally by monotonic, contextual, and periodic-like components, and expressed neurally through successive representational stages. More broadly, these results suggest that durations may be organised in a structured psychological space analogous, in principle, to perceptual spaces described in other domains. Characterising that space more fully may help bridge classical psychophysics, neural dynamics, and representational geometry, and provide a new framework for understanding how the mind and brain represent durations.

## Materials and Methods

### Study design

We combined a behavioural similarity-judgment task and an EEG duration-oddball task, performed by the same 33 participants in two separate sessions, to characterise the psychological and neural representational geometry of a fixed set of ten auditory durations (Fig. 1).

### Participants

Thirty-three healthy adults (mean age: 26.97 years; 19 female) were recruited through university mailing lists and the RISC’s participant pool (https://www.risc.cnrs.fr/). All participants reported normal or corrected-to-normal vision, and no history of neurological or psychiatric disorders. All were naive as to the purpose of the study.

### Ethics information

The study was approved by the Ethics Committee for Research (CER) of Paris-Saclay University (CER-Paris-Saclay-2023-089). Written informed consent were obtained from all participants in accordance with the guidelines of the Ethics Committee. Participants were compensated 20 euros for the first behavioural session and 90 euros for the second EEG session.

### Similarity judgment task (behaviour)

Both the similarity judgment and oddball detection tasks were implemented in PsychoPy v2021.2.3 (*86*). The experiment consisted of two separate sessions conducted in a quiet EEG-dedicated room at NeuroSpin (CEA/DRF, Gif-sur-Yvette, France), where participants remained seated comfortably in an armchair throughout the procedure.

To investigate the structure of duration representations, participants performed a similarity judgment task involving pairs of auditory (440 Hz pure tone) durations (Fig. 1). Each duration was drawn from a fixed set of ten durations ranging from 400 ms to 2200 ms in 200 ms steps (i.e., covering sub- to supra-second range). On each trial, pairs of durations were presented with an inter-stimulus interval of 700 ±10% ms. Their order was counter-balanced. To ensure that similarity judgments were based on duration rather than accumulated perceptual intensity, the loudness of each stimuli was randomly varied by ±5 dB around a base level of 70 dB, thereby reducing the potential influence of energy-based cues associated with longer durations (*87*). All 45 possible pairs of durations were presented in both orders, yielding a total of 90 pairs of durations per participant. Participants judged the perceived similarity of the pair on each trial (1 = "most dissimilar", 7 = "most similar") using the computer mouse. A 1000 ms inter-trial interval followed each response before the onset of the next trial.

Before the task, participants received standardised instructions both orally and in written form on screen, followed by a short practice phase consisting of 20 trials (duration pairs) composed of the most representative durations of the stimulus set (i.e., the smallest, the largest, and three intermediate durations), allowing them to familiarise themselves with the procedure and the rating scale. The task was divided into four blocks, each containing all 90 duration pairs presented in a randomised order. Each block lasted approximately 10 minutes, for a total session duration of approximately 50 minutes. Short breaks were offered between blocks. Participants could also take self-paced pauses within a block by withholding their response before initiating the next trial. Across the four blocks, each participant completed 360 trials, yielding a complete matrix of pairwise similarity ratings among the ten durations. Participants were explicitly instructed to refrain from counting or tapping to estimate durations (*88*).

### Oddball duration task (EEG)

Participants returned to the lab for the second session, approximately one week after the behavioural session. During this EEG session, participants performed a duration oddball detection task, designed to capture neural responses underlying the encoding of durations in the absence of explicit motor demands (i.e., button presses). Participants listened to a continuous stream of durations (as in the behavioural task, 440 Hz pure tones). Standard durations in the stream were randomly selected from the set of ten durations used in the similarity judgment task (400 ms to 2200 ms in 200 ms steps; 80% of trials). The deviant durations were 70 ms and 3400 ms (20% of trials) and were selected based on pilot data (N = 5) to ensure reliable perceptual discrimination from the standards. Participants were instructed to monitor the duration of each tone and detect oddballs (deviant durations) by pressing the space bar as quickly and as accurately as possible. The shorter- and longer-than-standard oddballs were included to ensure that participants monitored the full temporal extent of each stimulus. As in the similarity judgment task, stimulus intensity was randomly jittered by ±5 dB around a baseline of 70 dB to minimise reliance on cumulative intensity as a cue, ensuring that participants focused on temporal duration per se. The inter-stimulus interval was fixed at 700 ms, jittered by ±10% to prevent rhythmic entrainment (see Fig. 1 for a schematic representation of the trial sequence).

A trial-level feedback system was implemented to maintain task engagement. Correct detections (hits) were signalled by a green fixation cross, whereas missed oddballs or responses to standard stimuli (false alarms) triggered a red fixation cross. Only standard trials without response were retained for subsequent analyses.

### EEG data acquisition

EEG data were recorded using a 64-channel Waveguard cap (ANT Neuro, Enschede, Netherlands) mounted on the participant’s scalp. CPz served as the online reference electrode. EEG data were acquired using an eego mylab amplifier system at a sampling rate of 1000 Hz, a 131 Hz low-pass filter applied online and no high-pass filter. Impedances were kept below 20 kΩ at all times. Vertical eye movements were monitored via electrooculography (EOG), with one electrode placed above the left eye.

### EEG data preprocessing

Continuous EEG data were preprocessed using custom Python scripts and MNE-python software (*89*). Noisy or malfunctioning (e.g., flat) electrodes were identified by visual inspection, and participant-specific bad channels were annotated. Time periods containing muscular artifacts were also annotated. Ocular and other stereotyped artifacts were corrected using independent component analysis (ICA) computed separately for each block. ICA was fit on a temporally filtered copy of the data (high-pass filtered above 1 Hz, using MNE-Python’s default zero-phase FIR filter) to improve decomposition stability, while the resulting unmixing was applied to the corresponding unfiltered continuous signal. Components reflecting eye blinks and saccades were identified using EOG-channel correlations and marked for rejection, after which ICA was applied to obtain cleaned continuous data. The cleaned signal was then re-referenced to the common average reference, and previously marked bad channels were interpolated using spherical splines. Mastoid electrodes (M1 and M2) were excluded from all analyses, as they consistently exhibited high noise levels across participants and do not capture cortical activity (*90*). The cleaned continuous data were band-pass filtered (0.1-40 Hz, using MNE-Python’s default zero-phase FIR filter) and segmented into epochs time-locked to duration offset. At epoching, time segments previously annotated as artifacts were automatically excluded. Finally, we removed trials containing participants’ responses and trials corresponding to "odd" durations, before applying a baseline correction using a [-100, 0] ms interval relative to duration offset.

### Behavioural data analysis

#### Preprocessing similarity ratings

We summarised the similarity judgments of each participant in a representational dissimilarity matrix (RDM) according to the following procedure. Each similarity rating, initially provided on a 7-point Likert scale (1 = the most dissimilar, 7 = the most similar), was rescaled to the [0, 1] interval and converted to a dissimilarity score (i.e., 1 – rescaled similarity). For each participant, dissimilarity scores were then averaged across all repetitions of a given duration pair, yielding a 10 × 10 RDM in which each entry reflects the perceived dissimilarity between two durations.

#### Multidimensional scaling

To visualise the global structure of the duration space, we applied non-metric multidimensional scaling (MDS) (*14*) to each participant’s RDM. This technique embeds high-dimensional dissimilarity data into a low-dimensional geometric space while preserving the rank order of pairwise distances. To determine the dimensionality of the MDS solution, we computed stress values for embeddings spanning one to five dimensions. Stress quantifies the mismatch between the dissimilarities in the original RDM and the distances between points in the reduced-dimensional space, thereby providing a criterion for selecting the number of dimensions that best captures the structure of the data (*14, 16*).

#### Theoretical models

We next sought to assess how well the structure of the duration space, captured in the empirical RDM, could be explained by different theoretical accounts of duration representation. To this end, we constructed a set of theoretical RDMs, each embodying a specific hypothesis about the geometry of duration encoding (Table 1).

**Table 1.**
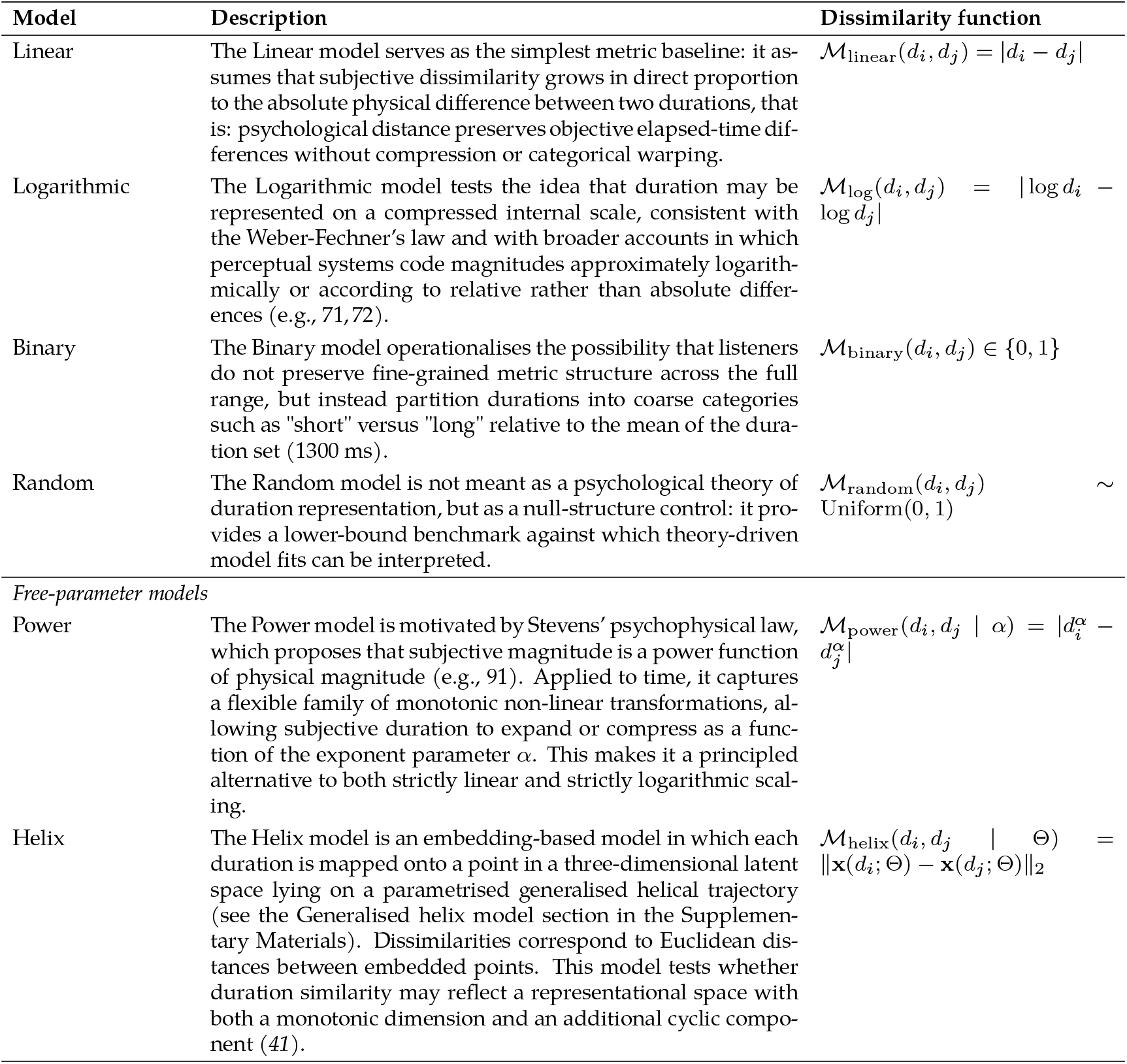
Summary of theoretical models and their dissimilarity functions. Free parameters of the Power and Helix models were optimised via an iterative optimisation process to find the configuration that maximised the Spearman correlation between the model-based RDM and the empirical RDM for each participant.

To assess how well each model accounted for participants’ empirical similarity judgments, we computed the Spearman rank-order correlation between the empirical RDM and each theoretical RDM. Before computing the correlation, we extracted the upper triangular part of each matrix (excluding the diagonal) to avoid redundancy and trivial self-comparisons. These elements were then flattened into 1D vectors of length 45. The resulting vectors captured the full set of unique pairwise dissimilarities in each RDM. The Spearman correlation coefficient thus provided a measure of model-data fit, robust to non-linear scaling. This approach allowed us to compare the internal structure predicted by each theoretical model to the structure derived from participants’ subjective similarity judgments.

#### Model comparison

Because theoretical RDMs vary in flexibility (0, 1, or 2 free parameters), we used a leave-one-duration-out cross-validation procedure to obtain an unbiased estimate of their predictive accuracy and prevent overfitting. For each left-out duration *d_k_* (10 folds in total):

1. **Training set.** We removed duration *d_k_* and its associated row and column from the empirical RDM, yielding a 9 × 9 sub-RDM. Parametrised models (i.e., Helix and Power) were fitted only to this sub-RDM, by minimising the negative Spearman correlation between the flattened upper-triangular part of the empirical and model-predicted RDMs.
2. **Testing set.** Using the fitted parameters 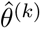, each model generated a full 10 × 10 predicted RDM. The model’s *out-of-sample prediction error* was computed exclusively on the left-out row and column (i.e., all dissimilarities involving *d_k_*).
3. **Fold score.** We quantified predictive accuracy for fold *k* as the Spearman correlation between the empirical dissimilarities *ε* (*d_k_, d_j_*) and model-predicted dissimilarities 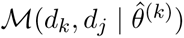 for all *j* ≠ *k*.

This produced 10 fold-wise correlation values per model:

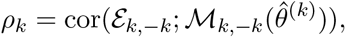

which we then averaged to yield the average cross-validated model score:

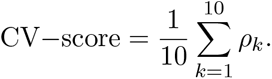

We stored the fitted parameters from each fold to assess parameter stability across folds. Models were compared on their median cross-validated correlation, which provides an unbiased estimate of how well each model predicts dissimilarities for durations it was *not* trained on. Because the metric reflects genuine out-of-sample performance, more flexible models only outperform simpler ones if their additional parameters capture structure that generalises across durations, rather than overfitting idiosyncrasies of the empirical RDM. After model comparison, we also fitted each parametrised model to the full empirical RDM (using the same correlation-based objective) to obtain a single set of interpretable parameters for visualisation and analysis. These "full-data" fits were not used for model comparison.

#### Group-level statistical inference

To assess the results of the model comparison procedure while accounting for the repeated cross-validation estimates obtained within participants, we fitted a Bayesian multilevel distributional model using the brms R package (*92, 93, 94*). Cross-validated correlation scores were first Fisher *z*-transformed using *z* = atanh(*r*). The model was fitted to these Fisher-transformed scores using a Student-*t* likelihood. Participant-level deviations were estimated for each theoretical model, thereby accounting for the repeated fold-level estimates within participants and for between-participant variability in model performance. In addition, the residual dispersion parameter was itself modeled as a function of theoretical model identity, with participant-level deviations, allowing the variability of CV scores to differ across models and participants. Weakly informative priors were used for all model parameters. Four Markov Chain Monte Carlo chains were run, each with 5000 iterations including 1000 warmup iterations. Posterior convergence was assessed by visual inspection of trace plots, the Gelman-Rubin statistic *R̂*, and effective sample sizes.

For inference on model comparisons, we did not use Bayes factors based on the Savage-Dickey density ratio, as these can be sensitive to prior specification for point-null hypotheses. Instead, we used the posterior distribution of pairwise differences between theoretical models. For each posterior draw, expected CV scores were obtained from the fitted model, back-transformed to the correlation scale using the inverse Fisher transformation, and averaged within each theoretical model. Pairwise differences between models were then computed at the draw level, yielding a posterior distribution for each contrast. These posterior contrasts were summarised using their posterior median and 95% highest density interval (HDI). To assess whether differences were practically meaningful, we additionally used a region of practical equivalence (ROPE) around zero, following the HDI+ROPE approach (*95, 96*). Evidence for a meaningful difference was assessed by examining whether the 95% HDI lay outside the ROPE, as well as by reporting the proportion of posterior mass falling inside the ROPE and on either side of zero. In the present analyses, the ROPE was defined as a difference in CV correlation between −0.01 and 0.01, corresponding to effects considered too small to be practically meaningful.

### EEG data analysis

#### Event-related potentials

In a first step, we aimed at identifying clusters of sensors that responded to (or "encoded") durations. To this end, we fitted a regression to predict EEG activity at duration offset for each participant, sensor, and timestep, using durations recoded using their rank (in ascending order) as a predictor. We used ranks not to impose a linear constraint on the identification of the channels, as this would bias the subsequent RSA in favour of the linear model. This analysis resulted in a slope estimate representing the monotonic effect of duration for each participant, sensor, and timestep. We then concatenated these 2D matrices into a 3D tensor of shape participants, channels, timesteps, and used a cluster-based permutation test with threshold-free cluster enhancement (TFCE) (*97*) to identify clusters of sensors and timesteps with slopes significantly different from 0.

#### Time-frequency analysis

To analyse induced oscillatory power across frequency bands at duration offset, we computed time-frequency representations (of the EEG temporal signals without baseline correction) using Morlet wavelets. These analyses were performed over a range spanning −2000 to 2000 ms relative to duration offset. Time-frequency decomposition was conducted across frequencies from 4 to 40 Hz in 1-Hz steps, encompassing canonical frequency bands from theta to low gamma. Frequencies below 4 Hz were excluded due to the limited epoch length. The number of wavelet cycles was set to half the center frequency, providing an optimal trade-off between temporal and spectral resolution. Time-frequency representations were then cropped to provide the final analysis window, spanning from −100 ms to 600 ms relative to duration offset. These power estimates were then normalised by taking the logarithm of the ratio of the period of interest over a [-100 ms, 0] baseline period (relative to offset). To isolate the induced oscillatory activity, we subtracted the evoked response (i.e., the across-trial per-condition average) from each epoch (*98*). We then averaged power in the theta (4–7 Hz), alpha (7–13 Hz), beta (13–30 Hz), and low-gamma (30–40 Hz) bands.

#### Representational similarity analysis

To characterise the relationship between similarity judgments, theoretical models, and EEG representations of duration in the oddball task, we performed a time-resolved representational similarity analysis (RSA). This analysis relied on the same set of theoretical RDMs used for the behavioural analyses, together with participant-level RDMs derived from EEG data. In addition to the theoretical models previously described (Table 1), we designed a novel RDM capturing similarities in the difference between each duration and the closest odd duration (i.e., 70 ms or 3400 ms), thus capturing a unique task-relevant feature.

EEG RDMs were computed within the [-100, 600] ms interval relative to duration offset using cross-validated representational dissimilarities, as implemented in the mne-rsa toolbox (*99*). For each time point, we estimated a 10×10 pairwise dissimilarity matrix using a cross-validated squared Euclidean distance (i.e., a cross-validated dissimilarity metric derived from squared Euclidean distances computed on multichannel EEG patterns). Specifically, trial-level EEG patterns were partitioned into folds and dissimilarities were computed in a cross-validated manner across folds, yielding a time-resolved series of condition-by-condition RDMs for each participant. These time-resolved EEG RDMs were then compared to the theoretical RDMs using Spearman correlations.

To further disentangle the unique contribution of each theoretical model in explaining the EEG variance, we then applied commonality analysis (e.g., 100, 101, 102) to quantify the relationship between behavioural and EEG RDMs while accounting for multiple theoretical RDMs. This variance-partitioning approach enables the decomposition of shared variance into components that are common to behaviour, EEG, and the theoretical models, while statistically controlling for variance uniquely attributable to the remaining models.

This analysis yielded, for each participant, a time-resolved estimate of the unique (partial) correlation between EEG RDMs and each theoretical RDM. To assess whether these time courses differed across theoretical models at the group level, we fitted a series of Bayesian generalised additive multilevel models using the brms R package (*92, 93, 94*). This modelling framework provides posterior odds for non-null effects at each time point while explicitly accounting for temporal dependencies as well as within- and between-participant variability. Importantly, this approach has been shown to yield more reliable and temporally precise estimates of the onset and offset of time-resolved M/EEG effects than cluster-based permutation tests (*103*).

#### Linking duration representations to neural dynamics

To elucidate the neural correlates of the representational geometry revealed by the behavioural data, we next sought to relate parameters of the theoretical models fitted to the behavioural RDMs to EEG features measured both i) during task performance (hereafter referred to as task-related EEG features) and ii) during out-of-task periods (hereafter referred to as endogenous EEG features).

As task-related EEG feature, we used the amplitude and the latency of the offset P2, computed over six clustered frontocentral electrodes: FC1, FCz, FC2, C1, Cz, and C2 (*45, 46, 47, 48, 49*). We additionally analysed the average theta, alpha, beta, and gamma power in the [0 − 200 ms] interval following duration offset. As endogenous EEG features, we used the intercept and exponent of the aperiodic (1/f) component, as well as the peak frequency and power of the alpha and beta bands (*104*), estimated during a movement-free 15-to-45 s period (exact duration varied across participants) of between-block resting state.

To assess the robustness of the resting-state brain-behaviour associations while accounting for the number of tested associations, we fitted a Bayesian multivariate regularised regression model. The behavioural model parameters (*β*, *α*, and *ω*) were entered as outcomes, and resting-state EEG features were entered as predictors, including alpha and beta power, alpha and beta peak frequency, aperiodic offset, and aperiodic exponent. All variables were standardised prior to modelling, so regression coefficients can be interpreted as standardised partial associations. Fold-to-fold variability of the behavioural model parameters was used to approximate uncertainty in the participant-level parameter estimates. The model was fitted with a Student-*t* likelihood to reduce sensitivity to potentially influential observations. All EEG-behaviour slopes were estimated jointly, with a regularising prior placed on the slope coefficients to shrink weak and noisy associations toward zero. Posterior distributions were summarised using medians, 95% credible intervals, the posterior probability of a positive association, and the proportion of posterior mass falling inside a region of practical equivalence (ROPE; [−0.01, 0.01] in standardised units).

## Acknowledgments

We thank Joao Barbosa and Simone Viganò for their early feedback on this manuscript.

## Author contributions

Conceptualization: C.G., L.N., V.v.W.

Methodology: C.G., L.N., V.v.W.

Software: C.G., L.N.

Validation: V.v.W.

Formal analysis: C.G., L.N.

Investigation: C.G., L.N., V.v.W.

Resources: V.v.W.

Data curation: C.G., L.N.

Writing, original draft: C.G., L.N.

Writing, review & editing: C.G., L.N., V.v.W.

Visualization: C.G., L.N.

Supervision: V.v.W.

Project administration: V.v.W.

Funding acquisition: V.v.W.

## Data, code, and materials availability

All anonymised behavioural data, EEG epochs, experimental materials, and analysis code are publicly available via the Open Science Framework: https://osf.io/vhcek/. Participant identifiers in the shared files run up to 38; the five missing identifiers correspond to scheduled participants who did not attend and provided no data (no participant was excluded from the analyses).

## Supplementary Materials

### Materials and Methods

#### Generalised helix model

In this embedding-based model, each duration *d* is mapped onto a point in a three-dimensional latent space lying on a parametrised helical trajectory. The geometry of the helix is defined by three components:

- **Radial growth:**

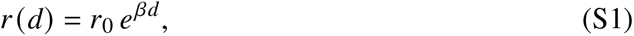

where *β* determines how the radius expands as a function of duration *d*.
- **Angular component:**

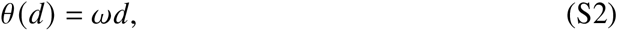

where *ω* sets the angular velocity (number of turns).
- **Vertical (z-axis) component:**

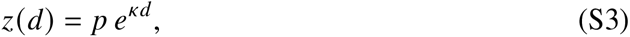

where *p* is the pitch (vertical spacing between turns) and *κ* determines exponential expansion along the vertical axis.

Combining these components, each duration is embedded in ℝ^3^ as:

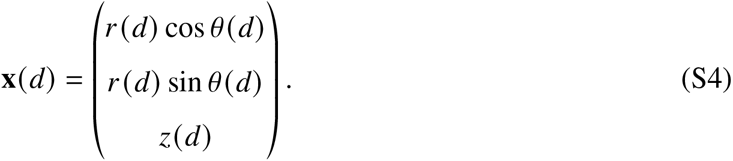

The predicted dissimilarity between two durations *d_i_* and *d _j_* is given by their Euclidean distance in this latent space:

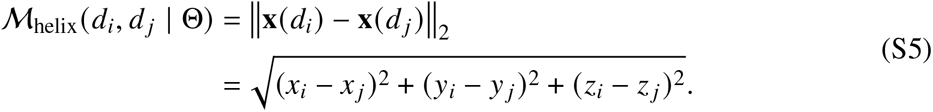

Here, Θ_helix_ = {*r*_0_, *β*, *p*, *ω*, *κ*} denotes the set of helix parameters. In the present study, a subset of these parameters was held constant (*r*_0_ = 0.1 and *p* = 1), *κ* and *β* were further assumed to be equal, and the remaining *two free parameters* (i.e., *β* and *ω*) were optimised separately for each participant by maximising the Spearman correlation between the model-predicted RDM and the empirical RDM. We provide a Shiny application allowing users to interact with the parameters of this model and visualise its predictions: https://lnalborczyk.github.io/apps/helix.

### Supplementary Text

#### Participant-level behavioural RDMs

Figure S1 shows the average symmetrised empirical RDM computed using the similarity judgements for each participant. As can be seen from this figure, participants showed a great amount of variability. Whereas all participants tended to show a clear diagonal pattern, and although these matrices contain the average of 2 orders × 4 repetitions = 8 similarity ratings for each pair of durations, participants varied in the precision of this pattern (i.e., dispersion around the diagonal), whether the subjective diagonal isochrony line coincided with the objective isochrony line (i.e., the matrix diagonal), or whether dispersion around the diagonal tended to increase or remain stable with duration.

**Figure S1:**
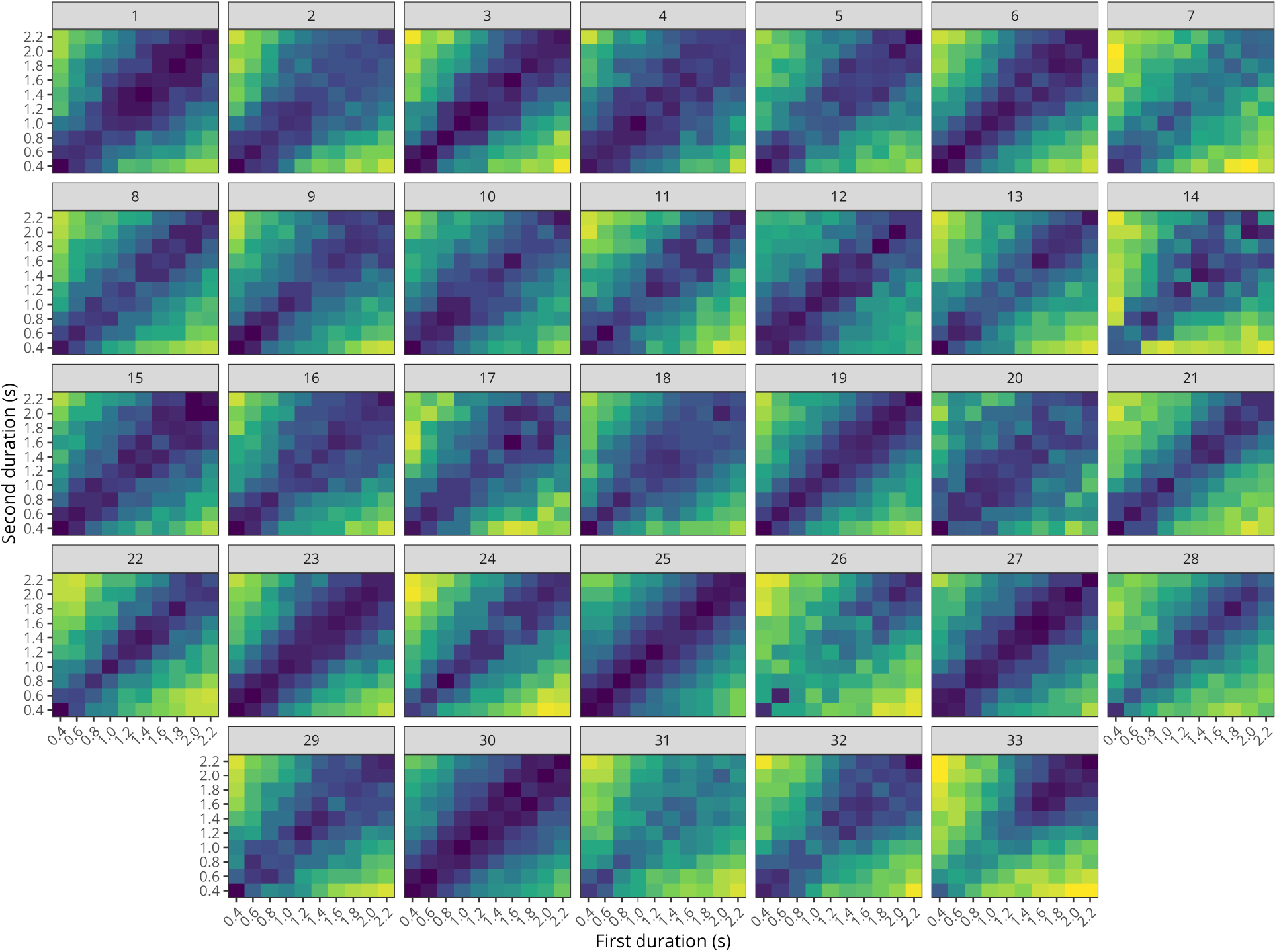
Participant-level average symmetrised behavioural RDMs. Each panel shows the average symmetrised empirical representational dissimilarity matrix for one participant.

#### Dimensionality of the behavioural duration space

To estimate the dimensionality of the behavioural duration space, we computed a scree function for each participant from their individual duration-based representational dissimilarity matrix. For each participant, non-metric multidimensional scaling was fit for solutions ranging from 1 to 5 dimensions, and the corresponding stress values were extracted. These participant-level stress values were then averaged across participants for each dimensionality, and the standard error of the mean was computed to characterise between-participant variability. To identify the elbow of the group-level scree plot, we applied two complementary heuristic criteria to the mean stress curve: (i) a maximum-distance method, which selects the dimensionality whose stress value lies farthest from the straight line connecting the 1- and 5-dimensional solutions, and (ii) a two-segment piecewise linear regression approach, which selects the breakpoint minimising the summed squared error across the two fitted segments. Both approaches converged on a 3-dimensional solution, which was therefore retained as the most parsimonious representation of the behavioural dissimilarity structure.

**Figure S2:**
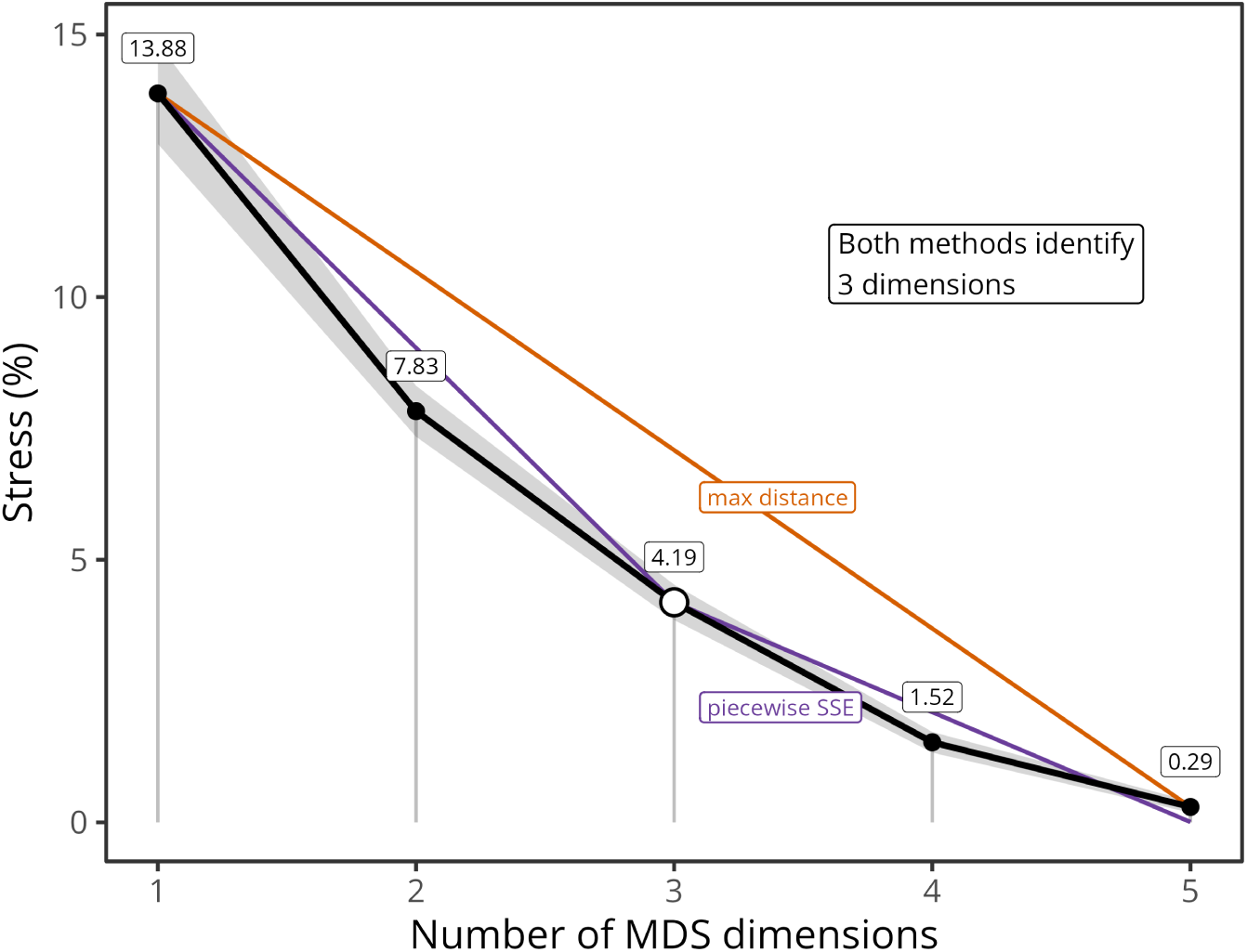
Group-level scree plot used to determine the dimensionality of the MDS solution. Mean stress values are shown for 1- to 5-dimensional non-metric MDS solutions, averaged across participants. The error band indicates the standard error of the mean. The black line and points show the group-average stress curve. The orange line corresponds to the chord used by the maximum-distance elbow criterion, and the purple line shows the fitted two-segment piecewise linear model. Both criteria identified an elbow at 3 dimensions, supporting retention of a three-dimensional MDS solution. Numeric labels indicate mean stress values at each dimensionality.

### Bayesian pairwise comparisons of behavioural RSA model performance

**Figure S3:**
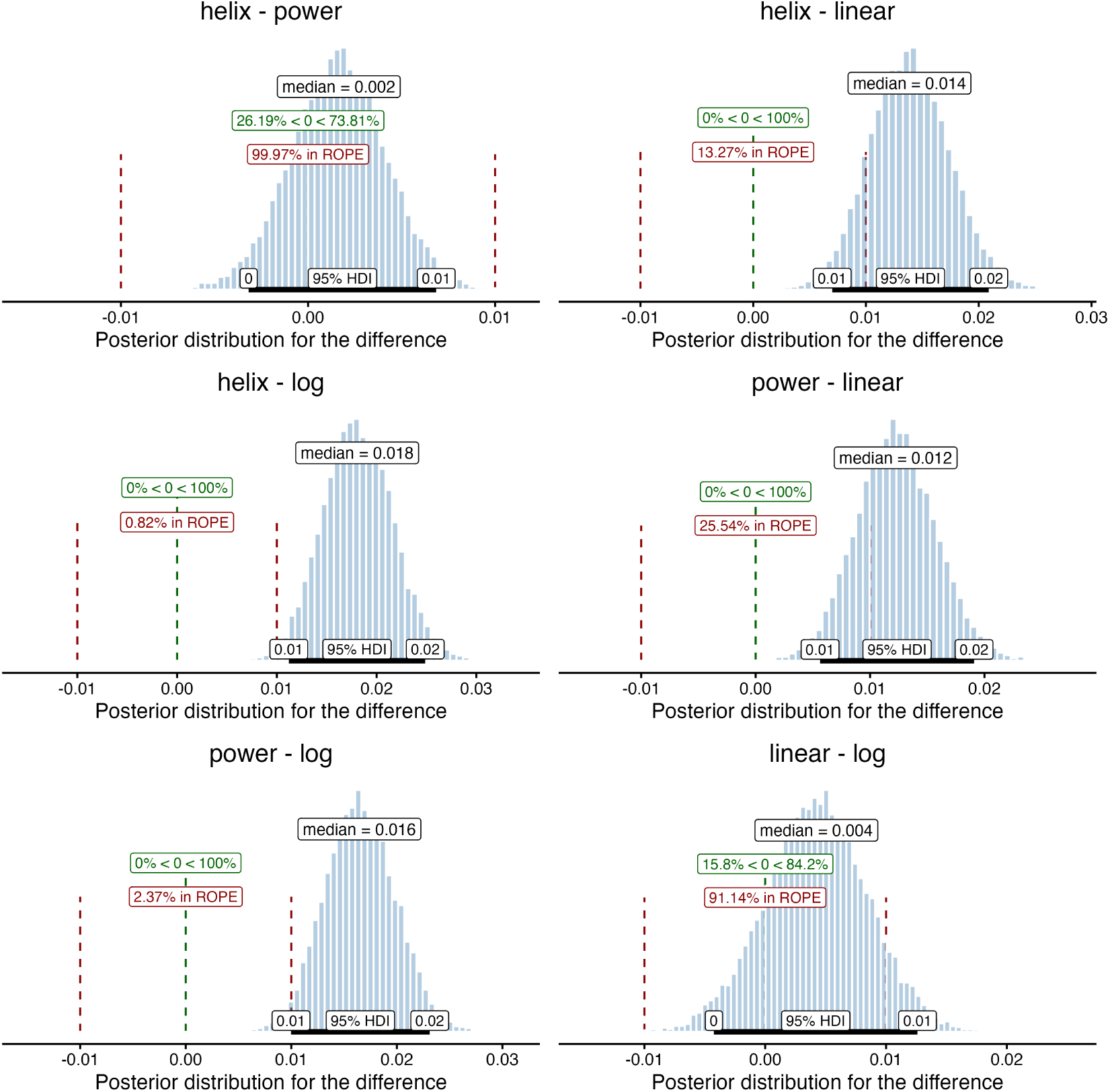
Posterior pairwise comparisons of cross-validated RSA model performance. Each panel shows the posterior distribution of the pairwise difference in expected cross-validated correlation between two candidate models, computed from the Bayesian multilevel model fitted to Fisher *z*-transformed CV correlations and back-transformed to the correlation scale. The black horizontal bar indicates the 95% highest density interval (HDI), the black label reports the posterior median, and the green dashed line marks zero. Red dashed vertical lines indicate the region of practical equivalence (ROPE; [−0.01, 0.01]). Green labels report the percentage of posterior mass below and above zero, whereas red labels report the percentage of posterior mass falling inside the ROPE.

### P2 amplitude and latency across durations

**Figure S4:**
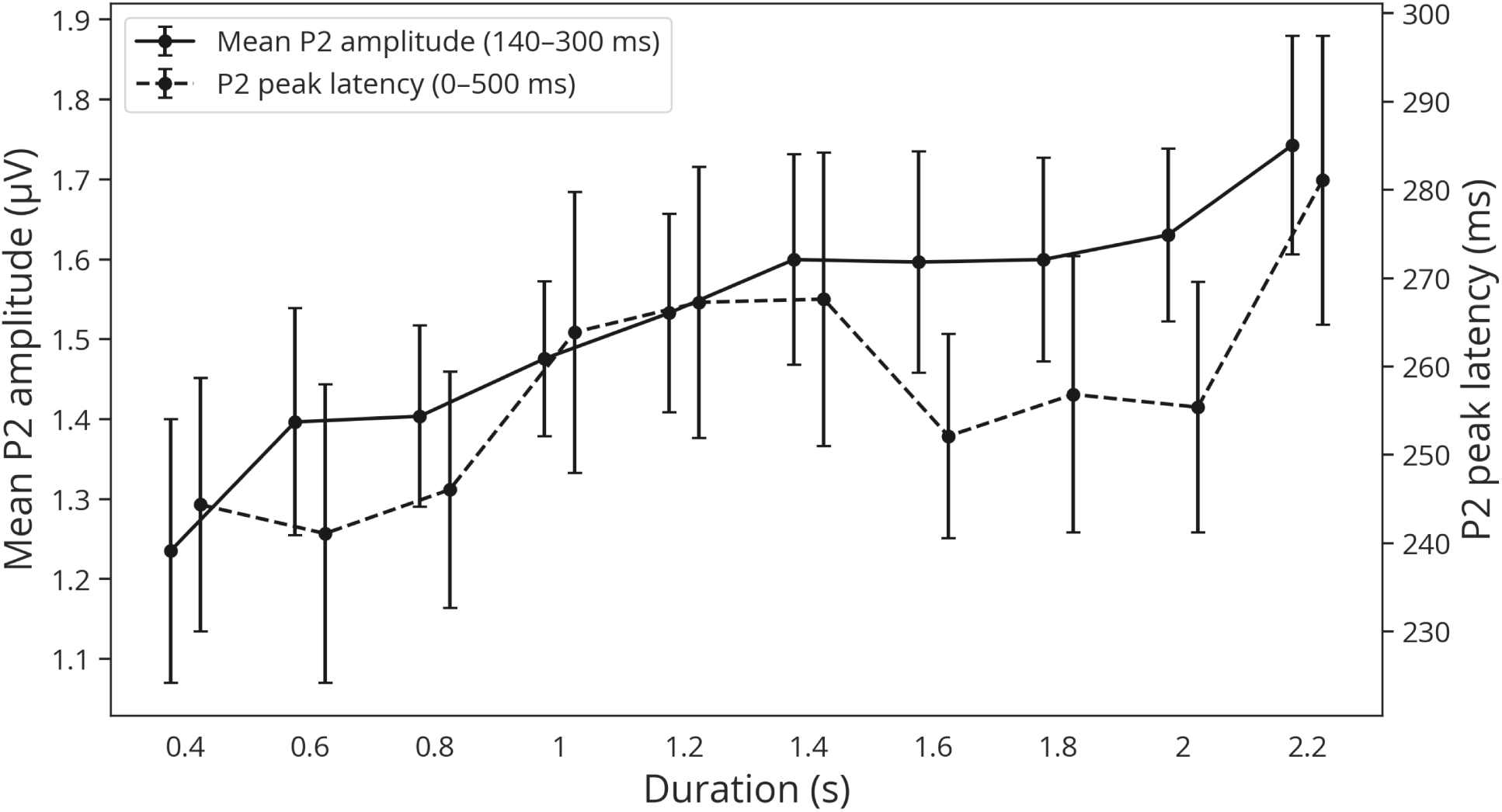
P2 amplitude and latency as a function of duration. Average P2 amplitude in the 140– 300-ms post-offset interval (left y-axis) and P2 peak latency (right y-axis). Dots represent group-level means and error bars represent the mean ± 1 SEM. While P2 amplitude scales monotonically with duration, P2 latency exhibits non-monotonic variations around intermediate and extreme durations.

